# The hydrogen sulfide donor sodium thiosulfate limits inflammation but aggravate smooth muscle cells apoptosis and aneurysm progression in a mouse model of abdominal aortic aneurysm

**DOI:** 10.1101/2023.09.15.557949

**Authors:** Clémence Bechelli, Diane Macabrey, Florian Caloz, Severine Urfer, Martine Lambelet, Florent Allagnat, Sébastien Déglise

## Abstract

**Intro:** The prevalence of abdominal aortic aneurysm (AAA) is constantly progressing with the aging of the global population. AAA rupture has a devastating 80% mortality rate and there is no treatment to slow-down AAA progression. Hydrogen sulfide (H_2_S) is a ubiquitous redox-modifying gasotransmitter produced in the cardiovascular system via the reverse trans-sulfuration pathway by cystathionine γ-lyase (CSE). H_2_S has protective properties on the cardiovascular system, including anti-inflammatory and antioxidant effects. Here, we hypothesized that sodium thiosulfate (STS), a clinically relevant source of H_2_S, would limit AAA growth.

**Methods:** 8-12 weeks old male WT or Cse^-/-^ mice on a C57BL/6J genetic background were submitted to a model of AAA by topical elastase application on the abdominal aorta and β-aminopropionitrile fumarate treatment in the drinking water for 2 weeks post-op. Sodium thiosulfate (STS) was given via the drinking water post-op until aorta collection. *In vitro* experiments were conducted to assess the effect of STS and pro-inflammatory cytokines interleukin-1 β and 6 and tumor necrosis factor α on primary human vascular smooth muscle cell (VSMC).

**Results:** Surprisingly, STS increased elastin degradation, AAA size and rupture, despite reducing infiltration of macrophages, antigen-presenting cells and lymphocytes in WT mice. Conversely, Cse^-/-^ mice with impaired H_2_S production developed smaller AAA than WT mice despite increased infiltration of immune cells. STS reduced VSMC coverage, possibly lowered VSMC proliferation, and promoted VSMC loss and extracellular matrix (ECM) breakdown. In vitro, STS aggravated pro-inflammatory cytokine-induced VSMCs apoptosis.

**Conclusion:** STS has a paradoxical effect on AAA growth, reducing inflammation while simultaneously impeding favorable vascular remodeling, resulting in bigger AAA in a model of periadventitial elastase. This study identifies a negative effect of H_2_S on VSMC in this environment, highlighting the complex role of H_2_S in AAA progression. The deleterious effect of STS on AAA progression is significant, especially given the growing use of STS in clinical settings for various indications.

## Introduction

Abdominal aorta Aneurysm (AAA) is a degenerative disease of the aorta wall that affects 5% of males aged 65 years^1, 2^. AAA is described as a local extension of the abdominal aorta wall that is greater than 50% larger than its usual diameter. AAA rupture has a devastating 80% death rate because most AAA are asymptomatic^3^. The only treatment options are open surgical repair or minimally invasive endovascular aortic repair (EVAR)^4^. Although the risk factors for AAA are widely documented (smoking, age, male sex, hypertension, atherosclerosis, genetic predispositions), the cellular and molecular mechanisms of AAA are not^5^. AAA’s primary pathogenic features are i) infiltration of innate and adaptive immune cells in the aorta wall; ii) loss of vascular smooth muscle cells (VSMCs); and iii) proteolysis of the extracellular matrix (ECM). The lack of resolution in those processes results in progressive AAA growth, culminating in AAA rupture. Overall, oxidative stress and inflammation are the primary causes of AAA^5, 6^. However, the precise sequence of events leading to rupture is unknown, and novel approaches to prevent aneurysm growth or rupture are required.

Hydrogen sulfide (H_2_S), a byproduct of the metabolism of sulfur-containing amino acids, is now commonly acknowledged as a gasotransmitter. H_2_S helps numerous organs and systems maintain homeostasis. H_2_S protects against vascular diseases through a variety of mechanisms, including the reduction of oxidative stress and inflammation, the improvement of EC function, NO production, and vasodilation, and the preservation of mitochondrial function^7,8^. H_2_S is produced in mammalian cells through the reverse transulfuration pathway by two pyridoxal 5ʹ-phosphate dependent enzymes, cystathionine γ-lyase (CSE) and cystathionine β-synthase^8^. Mice lacking Cse display may display age-dependent hypertension and have been shown to be adversely affected in various models of cardiovascular diseases^7^.

Endogenous H_2_S level and CTH expression are lower in AAA patients^9, 10^ and a few pre-clinical studies showed that H_2_S may provide benefits against aortic dissection^9, 11, 12^. Thus, NaHS attenuates inflammation and aortic remodeling in a model of aortic dissection induced by β-aminopropionitrile fumarate (BAPN) and angiotensin II (Ang-II) in WT mice^12^. Recently, aged Cse^-/-^ mice were shown to be more sensitive to angiotensin II-induced aortic elastolysis and medial degeneration, a phenotype rescued by NaHS treatment^9^. However, no study evaluated the role of CSE in AAA.

Given this evidence and the fact that H_2_S has anti-inflammatory and antioxidant properties^7, 13^, we hypothesized that sodium thiosulfate (Na_2_S_2_O_3_; STS), a clinically relevant source of H_2_S^14, 15^, might reduce inflammation and oxidative stress, thus limiting AAA progression. To test this hypothesis, we setup a mouse model of AAA using topical application of elastase on the sub renal aorta in WT or Cse^-/-^ treated with BAPN. The elastase AAA model is regarded as the best model for human AAA disease^16, 17^. Elastase breaks down medial elastin, leading to the formation of AAA^18, 19^. BAPN is a lysyl oxidase inhibitor that prevents cross-linking of elastin and collagen, leading to a chronic, growing AAA^19, 20^. In contrast to the results obtained with the AngII+BAPN model, no dissections are found in this model^20^. Surprisingly, STS facilitated AAA growth and induced rupture, whereas Cse^-/-^ mice were protected from AAA growth despite increased incidence of aortic dissection.

## Materials and methods

### Mice

WT mice C57BL/6JRj mice were purchased from Janvier Labs (Le Genest-Saint-Isle, France). Cse^-/-^ mice, kindly provided by Prof. James R Mitchell (Harvard T.H Chan School of Public Health, Boston, MA, USA), were bred and housed in our animal facility and genotyped as previously described^14^. All mice were housed at standard housing conditions (22°C, 12h light/dark cycle), with ad libitum access to water and regular diet (SAFE°150 SP-25 vegetal diet, SAFE diets, Augy, France). Mice were randomly treated or not with STS (Hänseler AG, Herisau, Switzerland) in the water bottle at 4g/L (0.5g/Kg/day), changed 3 times a week. BAPN (3-aminopropionitrile fumarate salt; SIGMA, A3134-5G) was dissolved in drinking water at 0.2% concentration and provided to the mice the day after the surgery until the end of the study.

Abdominal aorta aneurysm surgery was performed under isoflurane anesthesia (2.5% 2.5liter O_2_) as previously described^19, 20^. Analgesia was ensured by subcutaneous injection of buprenorphine (0.1 mg/kg Temgesic, Reckitt Benckiser AG, Switzerland) and local anesthesia via subcutaneous injection with a mix of lidocaine (6mg/kg) and bupivacaine (2.5mg/kg) along the incision line. 15 minutes post-injection and while deeply anesthetized, a midline incision was made, and the aorta separated from the surrounding fascia below the kidneys. A Whatmann paper impregnated with 8 µL of pancreas porcine elastase solution (MERCK, E1250-100mg) was applied on the surface of the aorta and left in place for 10 minutes. Following Whatmann removal, the peritoneum cavity was rinsed with warm saline, the abdomen closed with sutures and the skin closed with staples. Buprenorphine was provided before surgery, as well as a post-operative analgesic every 8h for 36 hours. The animals were monitored twice daily for signs of distress during recovery. Aortas were collected 14 days post-op by cervical dislocation and exsanguination under isoflurane anesthesia followed by PBS and 4% buffered formaldehyde perfusion., fixed in buffered formalin and included in paraffin for histology studies.

All animal experimentations confirmed to *The National Research Council:* Guide for the Care and Use of Laboratory Animals^21^. All animal care, surgery, and euthanasia procedures were approved by the CHUV and the Cantonal Veterinary Office (SCAV-EXPANIM, authorization number 3703).

### Histology

Abdominal aortas were fixed in 4% buffered formaldehyde for 24hours at 4°C, transferred to PBS solution, and subsequently embedded in paraffin, cut into 5μm sections, and stained using *Van Gieson Elastic Laminae* (VGEL) staining as previously described^22, 23^.

*Polychrome Herovici staining* was performed on paraffin sections as described ^24^. Young collagen is stained blue, while mature collagen is pink. Cytoplasm is counterstained yellow. Hematoxylin is used to counterstain nuclei blue to black.

*Calponin, CD3, CD8, CD86, MPO, F4/80, Ki67, Caspase3, IL-6, MMP9, SMA and CD206 Immunohistochemistry* was performed on paraffin sections. After rehydration and antigen retrieval (TRIS-EDTA buffer, pH 9, 17 min in a microwave at 500 watts), immunostaining was performed on abdominal aorta sections using the Rabbit specific HRP/DAB detection IHC detection kit (ab236469) according to manufacturer’s instructions. Slides were further counterstained with hematoxylin.

### Western blot

Mice aortas or human vein segments were flash-frozen in liquid nitrogen, grinded to power and resuspended in SDS lysis buffer (62.5 mM TRIS pH6,8, 5% SDS, 10 mM EDTA). Protein concentration was determined by DC protein assay. 10 to 20 µg of protein were loaded per well. Primary cells were washed once with ice-cold PBS and directly lysed with Laemmli buffer as previously described^23, 25^. Lysates were resolved by SDS-PAGE and transferred to a PVDF membrane Immobilon-P. Immunoblot analyses were performed as previously described^25^ using the antibodies described in supplementary table S1. Membranes were revealed using Immobilon Western Chemiluminescent HRP Substate in an Azure Biosystems 280 and analyzed using the Fiji (ImageJ 1.53t) software. Protein abundance was normalized to total protein using Pierce™ Reversible Protein Stain Kit for PVDF Membranes.

### Reverse transcription and quantitative polymerase chain reaction (RT-qPCR)

Frozen abdominal aortas were homogenized in TriPure^TM^ Isolation Reagent (Roche, Switzerland), and total RNA was extracted according to the manufacturer’s instructions. After RNA Reverse transcription (Prime Script RT reagent, Takara), cDNA levels were measured by qPCR Powerup SYBR^TM^ Green Master Mix (Ref: A25742) in a QuantStudio 5 Real-Time PCR system (Applied Biosystems, Thermo Fischer Scientific, AG, Switzerland). We looked at the expression of PGC1α using 5’-TGCTGTGTGTCAGAGTGGATT-3’ as forward primer and 5’-AGCAGCACACTCTATGTCACTC-3’ as reverse primer.

### Proteomics analysis

*Sample preparation*: Flash frozen abdominal aortas were pulverized in liquid nitrogen, resuspended in lysis buffer, sonicated, and boiled. Samples were diluted 1:1 with triethylammonium bicarbonate buffer, digested by adding 0.1 µg of modified trypsin (Promega) and incubated overnight at 37 °C, followed by a second digestion for 2 h with the same amount of enzyme. The supernatant was collected, diluted twice with 0.1% formic acid and desalted on strong cation-exchange micro-tips (StageTips, Thermo Fisher Scientific) as described previously60. Peptides were eluted with 1.0 M ammonium acetate (100 µl). Dried samples were resuspended in 25 µl of 0.1% formic acid, 2% acetonitrile prior being subjected to nano liquid chromatography tandem mass spectrometry (LC–MS/MS).

### LC–MS/MS analysis

Tryptic peptide mixtures (5 µl) were injected on a Dionex RSLC 3000 nanoHPLC system interfaced via a nanospray source to a high-resolution QExactive Plus mass spectrometer (Thermo Fisher Scientific). Peptides were separated on an Easy Spray C18 PepMap nanocolumn (25 or 50 cm×75 µm ID, 2 µm, 100 Å, Dionex) using a 35 min gradient from 4 to 76% acetonitrile in 0.1% formic acid for peptide separation (total time: 65 min). Full mass spectrometry (MS) survey scans were performed at 70,000 resolutions. In data-dependent acquisition controlled by Xcalibur v.4.0.27.19 software (Thermo Fisher Scientific), the ten most intense multiply charged precursor ions detected in the full MS survey scan were selected for higher energy collision-induced dissociation (normalized collision energy= 27%) and analysis in the orbitrap at 17,500 resolution. The window for precursor isolation was of 1.6 m/z units around the precursor and selected fragments were excluded for 60 s from further analysis. Sample quality control was performed by label-free test on short gradients and analyzed using the MaxQuant software (version 1.5.3.3). Then, 6-plex TMT labelling was performed and samples were injected separately. MS data were analyzed using Mascot v.2.5 (Matrix Science) set up to search the UniProt (www.uniprot.org) protein sequence database restricted to *mus. musculus*. Trypsin (cleavage at K,R) was used as the enzyme definition, allowing two missed cleavages. Mascot was searched with a parent ion tolerance of 10 ppm and a fragment ion mass tolerance of 0.02 Da (QExactive Plus). Iodoacetamide derivative of cysteine was specified in Mascot as a fixed modification. N-terminal acetylation of protein, oxidation of methionine and phosphorylation of Ser, Thr, Tyr and His were specified as variable modifications.

### Computational analysis

Annotated raw counts were further filtered to keep proteins detected in at least 6 out of 8 samples and to remove duplicates. The raw counts were converted into counts per million (CPM) and protein expression was log 2 normalized. T-distributed stochastic neighbor embedding (t-SNE) was computed using the top 1000 most varying proteins, then reduced to 50 PCA dimensions before computing the t-SNE embedding. The perplexity heuristically set to 25% of the sample size or 30 at maximum, and 2 at minimum. Calculation was performed using the Rtsne R package.

For identification of differentially expressed proteins, multi-method statistical testing was employed ^26, 27^ using two independent statistical methods: DESeq2 (Wald test) ^28^ and edgeR (LRT test) ^29^. Proteins with a Log FC superior to 0.2 and FDR below 0.2 were considered significant. The maximum q-value of the two methods was taken as aggregate q-value, which corresponds to taking the intersection of significant proteins from the two tests.

Statistical testing of differential enrichment of protein sets was performed using an aggregation of multiple statistical methods: fGSEA, GSVA/limma, and ssGSEA. The maximum q-value of the selected methods was taken as aggregate meta.q value, which corresponds to taking the intersection of significant proteins from all tests. The enrichment score of a GO term was defined as the sum of q-weighted average fold-changes, (1-q)*logFC, of the GO term and all its higher order terms along the shortest path to the root in the GO graph. The fold-change of a protein set was defined as the average of the fold-change values of its members. This graph-weighted enrichment score thus reflects the enrichment of a GO term with evidence that is corroborated by its parents in the GO graph and therefore provides a more robust estimate of enrichment. Data preprocessing was performed using bespoke scripts using R (R Core Team 2013)^30^ and packages from Bioconductor^31^. Statistical computation and visualization have been performed using the Omics Playground version v3.2.25-master230905^32^.

### Primary human VSMC culture

Human VSMCs were prepared from human saphenous vein segments as previously described ^23, 25^. Vein explants were plated on the dry surface of a cell culture plate coated with 1% Gelatine type B (Sigma-Aldrich). Explants were maintained in RPMI, 10% FBS medium in a cell culture incubator at 37°C, 5% CO_2_, 5% O_2_ environment. 9 different veins/patients were used in this study to generate VSMC.

### RNA interference

CSE knockdown was performed using human siRNA targeting CTH (Ambion-Life Technologies, ID: s3710 and s3712). The control siRNA (siCtrl) was the AllStars Negative Control siRNA (Qiagen, SI03650318). VSMC grown at 70% confluence were transfected overnight with 30 nM siRNA using lipofectamin RNAiMax (Invitrogen, 13778-075). After washing, cells were maintained in full media for 48h prior to assessment.

### Mitochondrial network analyzes

The mitochondrial network was observed by live cell imaging using the Mitotracker Red CM-H_2_XRos fluorescent probe (Thermofischer, M7513). The probe was dissolved in anhydrous DMF at 1 mM and used at 1 μM in serum-free RPMI. Live-cell image acquisition was performed using a Nikon Ti2 spinning disk confocal microscope. Images were analyzed automatically using the MiNA (Mitochondrial Network Analysis) toolset^33^ in the Fiji (ImageJ 1.53t) software.

### Apoptosis and caspase 3/7 activity

VSMC were grown on a 96 well plate. The percentage of apoptotic cells was determined using the DNA-binding dyes propidium iodide (PI, 5 μg/ml) and Hoechst 33342 (HO, 5 μg/ml, Sigma-Aldrich) as previously described. The cells were examined by inverted fluorescence microscopy (Leica). A minimum of 500 cells was counted in each experimental condition by two independent observers, one of them unaware of sample identity. caspase 3/7 activity was measured using Apo-ONE® Homogenous Caspase 3/7 Assay (Promega). VSMC were grown on a 96 well plate and 50 µl of the reagent was added to 50 µl of medium in each well. Blank are composed of the reagent and medium. After one hour, fluorescence with an excitation’s wavelength of 485 ± 20 nm and an emission’s wavelength of 530±20 nm is detected in a multimode plate reader (Synergy H1, Biotek AG).

### Statistical analyzes

All experiments adhered to the ARRIVE guidelines and followed strict randomization. All experiments and data analysis were conducted in a blind manner using coded tags rather than actual group name. A power analysis was performed prior to the study to estimate sample-size. Based on previous experience, using a detectable difference of 40% in aorta diameter by histomorphometry, a standard deviation of 20%, a desired power (1-β) of 0.8, and p value of 0.05 (alpha =0.05), it was determined that a total of 10-12 animals in each group is necessary to reach statistically meaningful conclusions. All experiments were analyzed using GraphPad Prism 9. Normal distribution of the data was assessed using Kolmogorov-Smirnov tests. All data with normal distribution were analyzed by unpaired bilateral Student’s t-tests or Mixed-effects model (REML) followed by post-hoc t-tests with the appropriate correction for multiple comparisons. For non-normal distributed data, Kruskal-Wallis non-parametric ranking tests were performed, followed by Dunn’s multiple comparisons test to calculate adjusted p values. Unless otherwise specified, p-values are reported according to the APA 7^th^ edition statistical guidelines. *p<.05, **p<.01, ***p<0.001.

## Results

### STS promotes aneurysm growth in WT mice in the model of elastase-induced AAA

We first assessed whether STS treatment protected against AAA growth and rupture following surgery mouse surgery. Surprisingly, STS treatment (4g/L) increased elastolysis in WT mice (Fig. 1A, B) and reduced survival (Fig. 1C). STS also increased AAA size (Fig. 1A), as quantified as the area under the curve of the aortic lumen area over 2mm (Fig. 1D) and max lumen area diameter (Fig. 1E). CSE is the main enzyme responsible for endogenous H_2_S production in the vascular system and Cse^-/-^ mice have impaired H_2_S production capacity ^14^. Interestingly, Cse^-/-^ mice developed smaller AAA than their WT littermates (Cse^+/+^) (Fig. 1F-J) with comparable survival rates (Fig. 1H), despite increased incidence of elastin breaks (Fig. S1).

**Figure 1.**
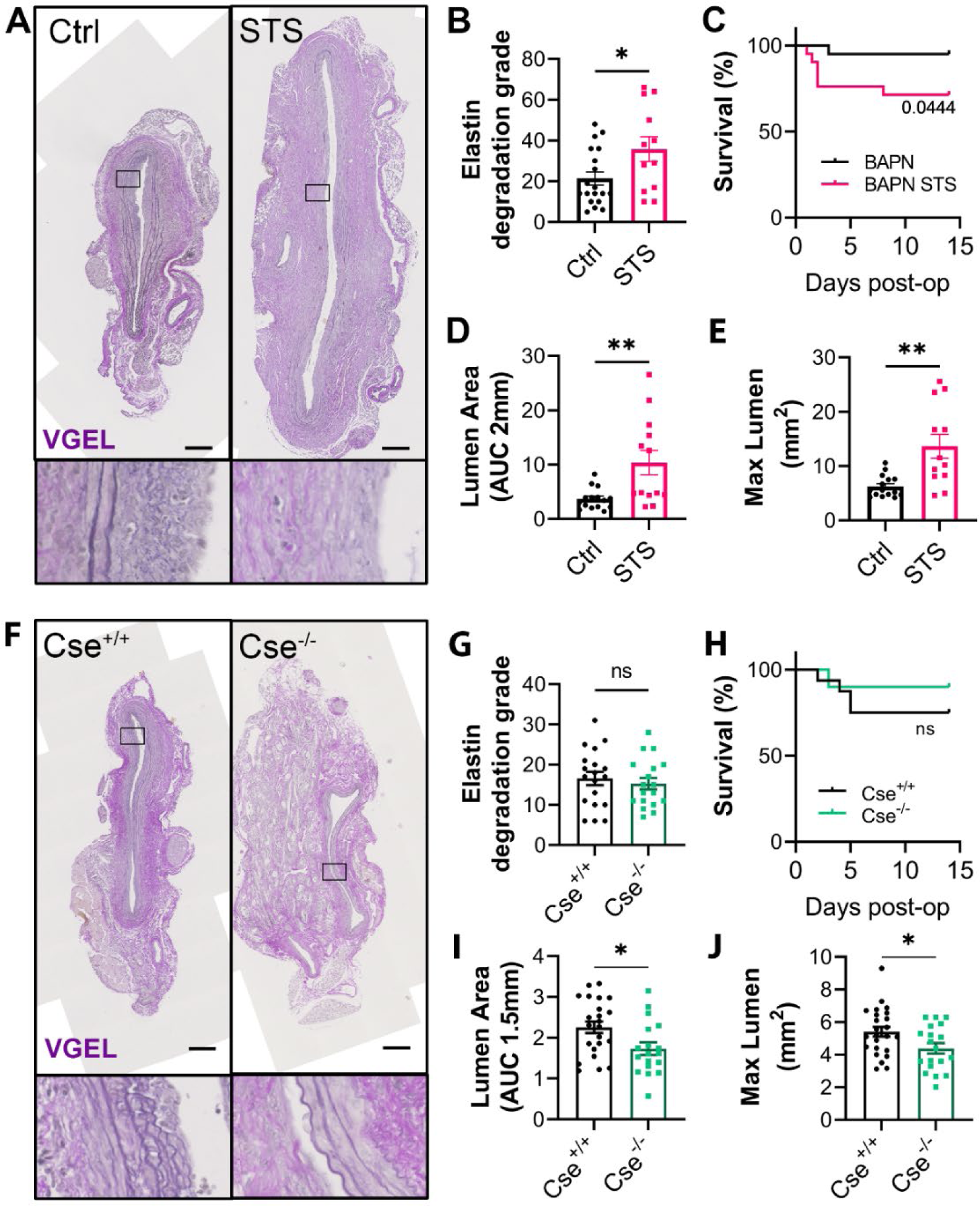
STS increases AAA size in a mouse model of topical application of elastase. Representative VGEL staining (A), elastin degradation grade (B), survival (C) quantitative assessment of aorta lumen area AUC over 2mm (D) and max lumen area (E) in sub-renal mouse aorta in WT mice with topical elastase application, treated or not (Ctrl) with 4g/L STS (STS). Representative VGEL staining (F), elastin degradation grade (G), survival (H), and quantitative assessment of aorta lumen area AUC over 1.5 mm (I) and max lumen area (J) in sub-renal mouse aorta in Cse^+/+^ or Cse^-/-^ with topical elastase application. (A, F) Scale bars = 100 μm. Lower Insets are 5-fold magnifications of main images. Data are mean±SEM of 15 to 18 animals per group. *p<0.05, **p<0.01 as determined by bilateral unpaired t-test.

### STS reduces inflammation in the aortic wall in the model of elastase-induced AAA

Inflammatory cells play a major role in the expansion of AAA ^6^. To study the impact of STS on inflammation, we measured immune cells infiltration in the AAA wall by histology. STS limited the infiltration of F4/80^+^ and CD206^+^ macrophages and CD86^+^ antigen-presenting cells (Fig. 2A, Fig. S2A) but not MPO^+^ neutrophils (Fig. S2B). STS also reduced the infiltration of CD3^+^, CD8^+^ and CD4^+^ lymphocytes (Fig. 2B). Similar investigation on Cse^-/-^ mice revealed an opposite impact on inflammation, with an increased infiltration of F4/80^+^macrophages, CD86^+^ antigen-presenting cells, and CD3^+^ and CD8^+^ lymphocytes (Fig. 2C-D), but not CD4^+^ cells and MPO^+^ neutrophils (Fig. S2C).

**Figure 2.**
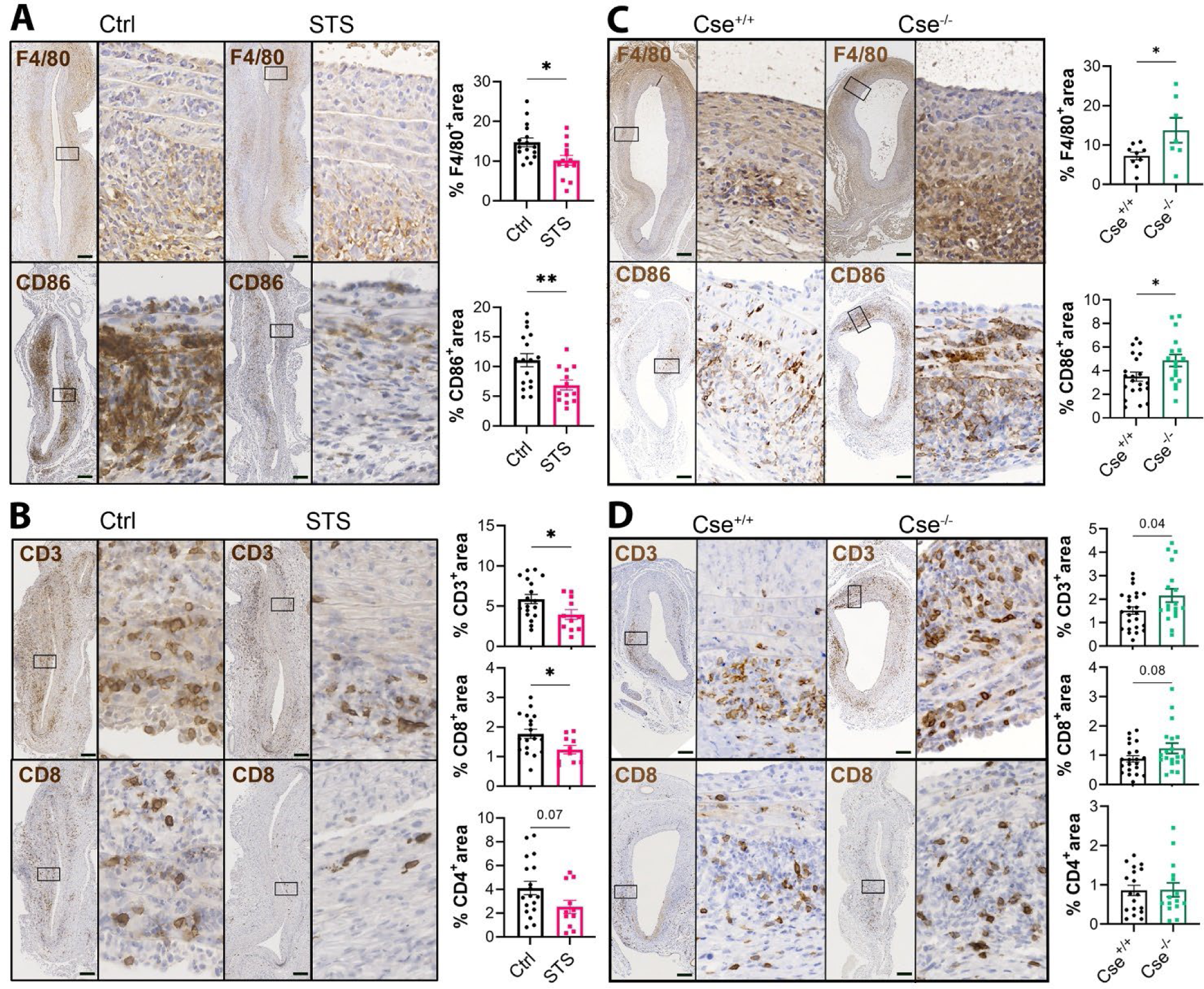
H_2_S reduces immune cells infiltration in the AAA wall. Representative F4/80, CD86 (A), CD3 and CD8 (B) immunostaining in sub-renal mouse aorta in WT mice with topical elastase application, treated or not (Ctrl) with 4g/L STS (STS). Representative F4/80, CD86 (C), CD3 and CD8 (D) immunostaining in sub-renal mouse aorta in Cse^+/+^ or in Cse^-/-^ mice with topical elastase application. Scale bars=100 μm. Right Insets are 5-fold magnifications of left images. Data are mean±SEM of 12 to 18 animals per group. *p<0.05, **p<0.01 as determined by bilateral unpaired t-test.

### STS increases mitochondrial biogenesis but negatively impact matrix remodeling

To determine the mechanism whereby STS impacted AAA growth despite anti-inflammatory effects, an untargeted approach was employed. Proteomic analysis of native aorta from WT mice treated or not with 4g/L STS for 1 week (n=4 per group) identified 2245 proteins (supplementary table S2), among which 119 up-regulated (LogFC=0.5, q=0.1; supplementary table S3), and only 4 down-regulated (LogFC =0.5, q=0.1; Fig. 3A-B, supplementary table S4). Pathway analysis revealed that STS selectively up-regulated processes linked to the mitochondria, including the electron transport chain, TCA cycle, and fatty acid beta-oxidation (Fig. 3C and table S5). Pathway analysis of down-regulated proteins revealed an enrichment of pathways associated with ECM remodeling (GO:0030198: extracellular matrix organization; FDR= .001) (Fig. 3C and table S5).

**Figure 3.**
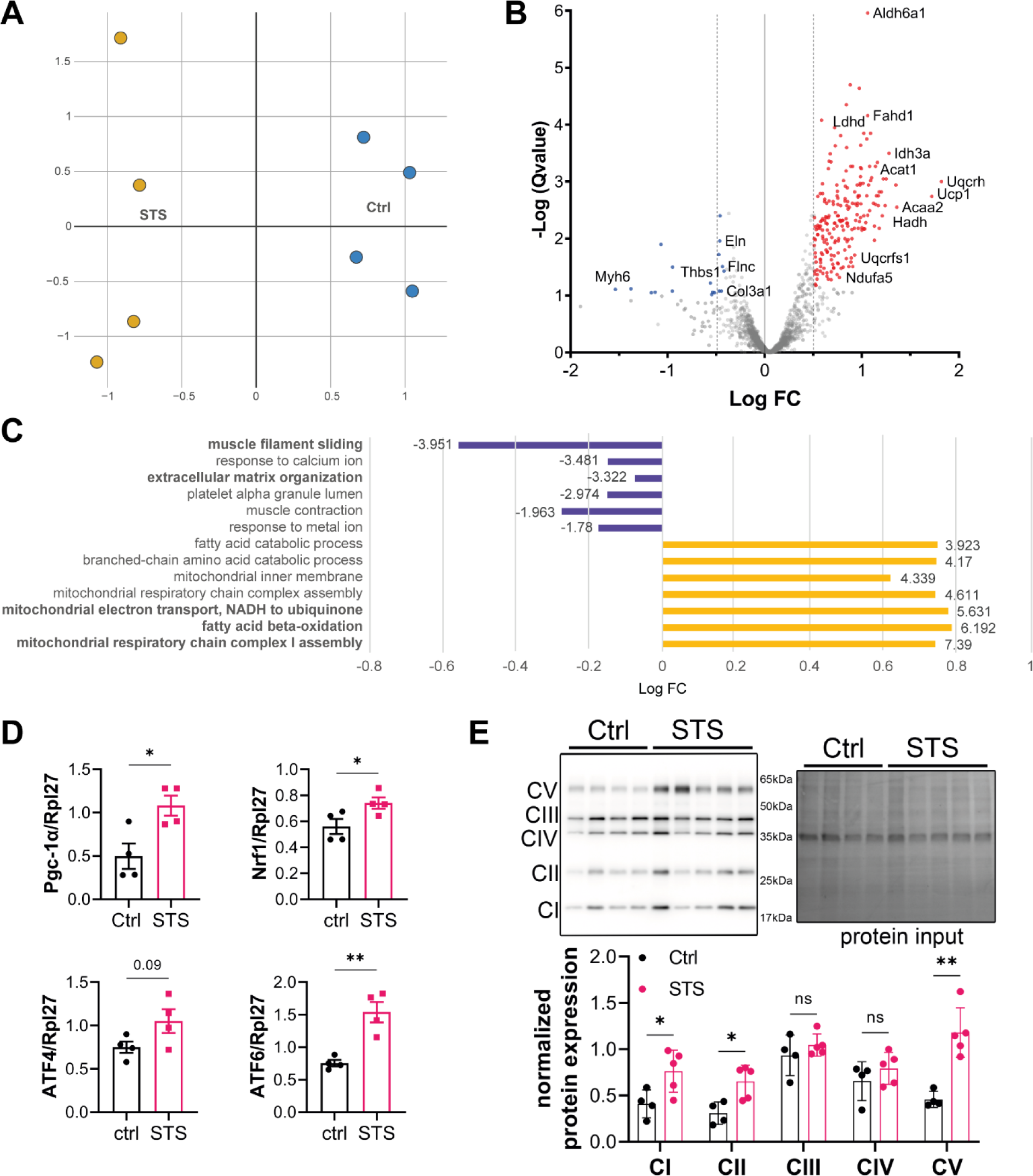
STS promotes mitochondrial biogenesis in the native aorta treated 1 week with STS. A) Principal component analysis of proteomics analysis from native aorta of WT mice treated or not (Ctrl) with STS (4g/L) for 1 week (4 animals per condition). B) Volcano plot showing differential protein expression (upregulated in red; down-regulated in blue) in Ctrl versus STS-treated aorta. C) Significantly down-regulated (blue) or up-regulated (yellow) gene ontology terms from pathway analysis expressed as Log fold change (FC). Numbers next to bars refer to enrichment score. D) qPCR analysis of mRNA expression in native aorta of WT mice treated or not (Ctrl) with STS (4g/L) for 1 week. Data are mean±SEM of 4 animals per group. *p<0.05 as determined by bilateral unpaired t-test. E) Representative Western blot of mitochondrial chain complex over total protein in native aorta of WT mice treated or not (Ctrl) with STS (4g/L) for 1 week. Data are mean±SEM of 5 animals per group.

The proteomic analysis uncovered uniform overexpression of mitochondrial proteins (Fig. 3C). H_2_S has also been proposed to rewire metabolism^34^ and promote mitochondrial biogenesis^35^. A one-week STS treatment increased the mRNA expression of the peroxisome proliferator-activated receptor gamma coactivator 1-alpha (PGC-1α), a master regulator of mitochondrial biogenesis^36^and nuclear respiratory factor-1 (NRF-1), as well as adaptive stress response transcription factors ATF4 and 6 in the abdominal aorta (Fig. 3D). It also increased the protein expression of complex I, II and V of the mitochondrial chain (Fig. 3E).

Among the mildly but significantly down-regulated proteins were Elastin (Eln), the main component of elastic laminae in the aorta, several collagen isoforms (Col4a6, Col5a1), and periostin (Postn), thrombospondin (Thbs1), and Filamin C (Flnc), proteins involved in ECM remodeling (Fig. 4A). In addition, STS treatment increased matrix metalloproteinase 9 (MMP9) in the AAA wall, a protease that degrade type IV and V collagens involved AAA^37, 38^ (Fig. 4B). Conversely, the MMP9 staining was reduced in Cse^-/-^ compared to Cse^+/+^ (Fig. 4C).

**Figure 4.**
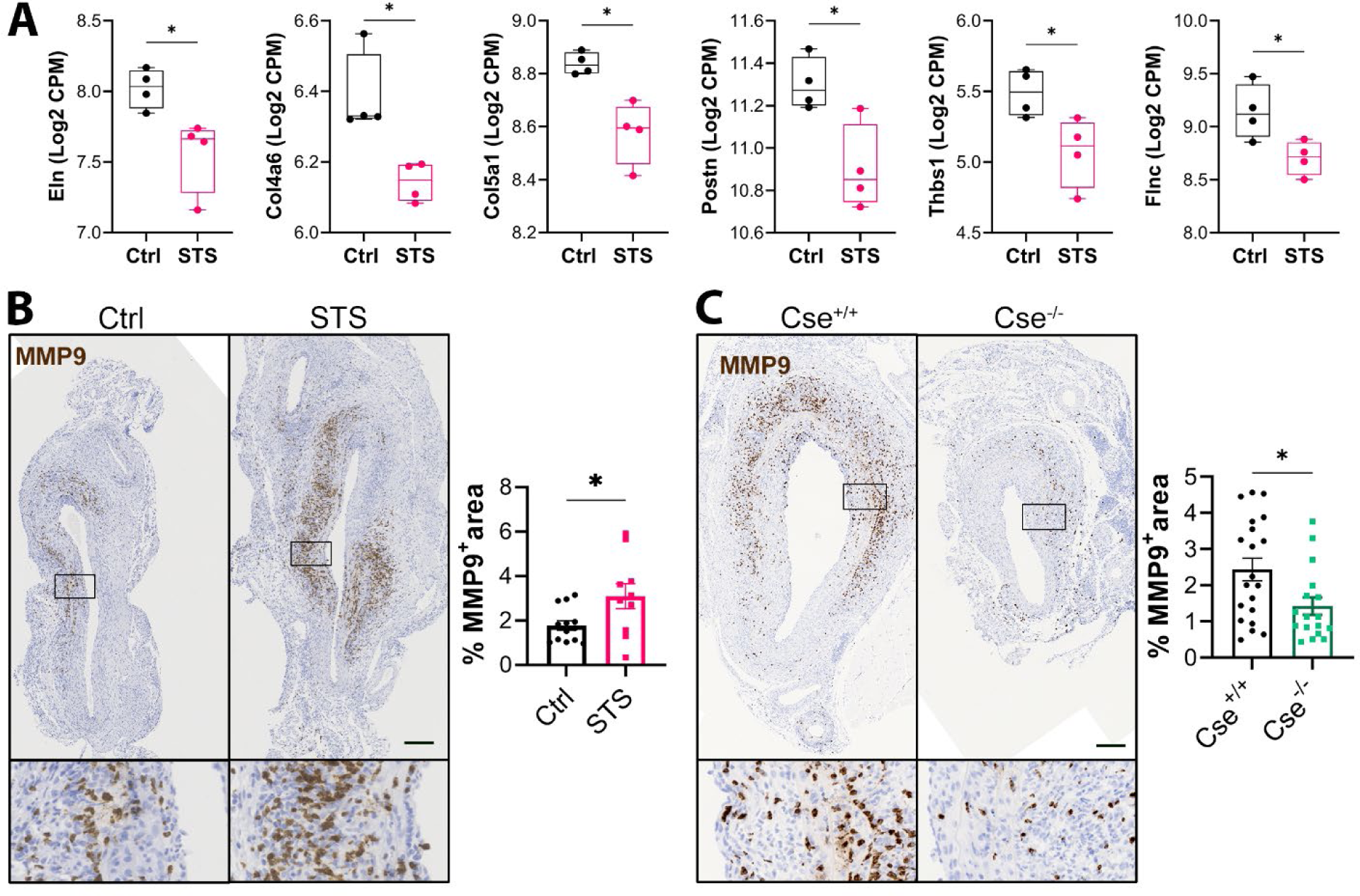
STS decreases extracellular matrix protein expression and promotes matrix degradation. A) Selected proteomics data (Log2 count per million (CPM) expression of individual proteins) in native aorta from WT mice treated ot not for 1 week with STS. *q<0.05 from proteomic analysis. Representative MMP9 immunostaining in sub-renal mouse aorta in WT mice treated or not (Ctrl) with STS (STS) (B), or in CSE^+/+^ or CSE^-/-^ mice (C). Lower insets are 5-fold magnifications of main images. Data are mean±SEM of 10 to 18 animals per group. Scale bar 100 μm. *p<0.05 as determined by bilateral unpaired t-test.

### STS increases cytokine-induced mitochondrial dysfunction and VSMC apoptosis

Surprisingly, when assessing de novo collagen deposition using polychrome Herovici staining, we observed increased collagen deposition in STS-treated mice (Fig. 5A) despite reduced collagen expression (Fig. 4A-B). This could suggest fewer cells, which could be linked to reduced inflammation (Fig. 2) or reduced compensatory VSMC expansion, which occurs during AAA formation^37^. Of note, we previously showed that STS inhibits VSMC proliferation in the context of intimal hyperplasia^14^. However, cell proliferation, as assessed by Ki67 staining, did not reveal any effect of STS on proliferation (Fig. 5B). We further evaluated the VSMC phenotype in AAA using the marker of contractile VSMC Calponin. STS decreased the number of Calponin^+^ cells (Fig. 5C), although it did not impact VSMC phenotype (calponin, SMA and SM22α expression) in native aortas (Fig. S3A). There was no difference in Calponin^+^ cells in Cse^-/-^ mice (Fig. 5D). VSMC apoptosis is a known feature of AAA progression and rupture ^37^. We could not evaluate the impact of STS on apoptosis *in vivo* due to very low numbers of cleaved caspase 3^+^ cells in AAA samples (Fig. S3B). To mimic the pro-inflammatory environment of AAA, we treated primary human VSMC with a cocktail of pro-inflammatory cytokines composed of IL-1β, IL-6, and TNFα, which are prominent in AAA^39, 40^. STS alone did not induce apoptosis but promoted cytokine-induced apoptosis (Fig. 6A) and cleaved caspase 3/7 activity (Fig. 6A). In contrast, siRNA-mediated CTH knock-down (Fig S4) protected against cytokine-induced VSMC apoptosis (Fig. 6B). STS also aggravated the impact of cytokines on the Bax/Bcl2 ratio (Fig. 6C) in VSMC and worsen cytokine-induced mitochondrial dysfunction as assessed by live Mitotracker staining and determination of the mitochondrial network (Fig. 6D).

**Figure 5.**
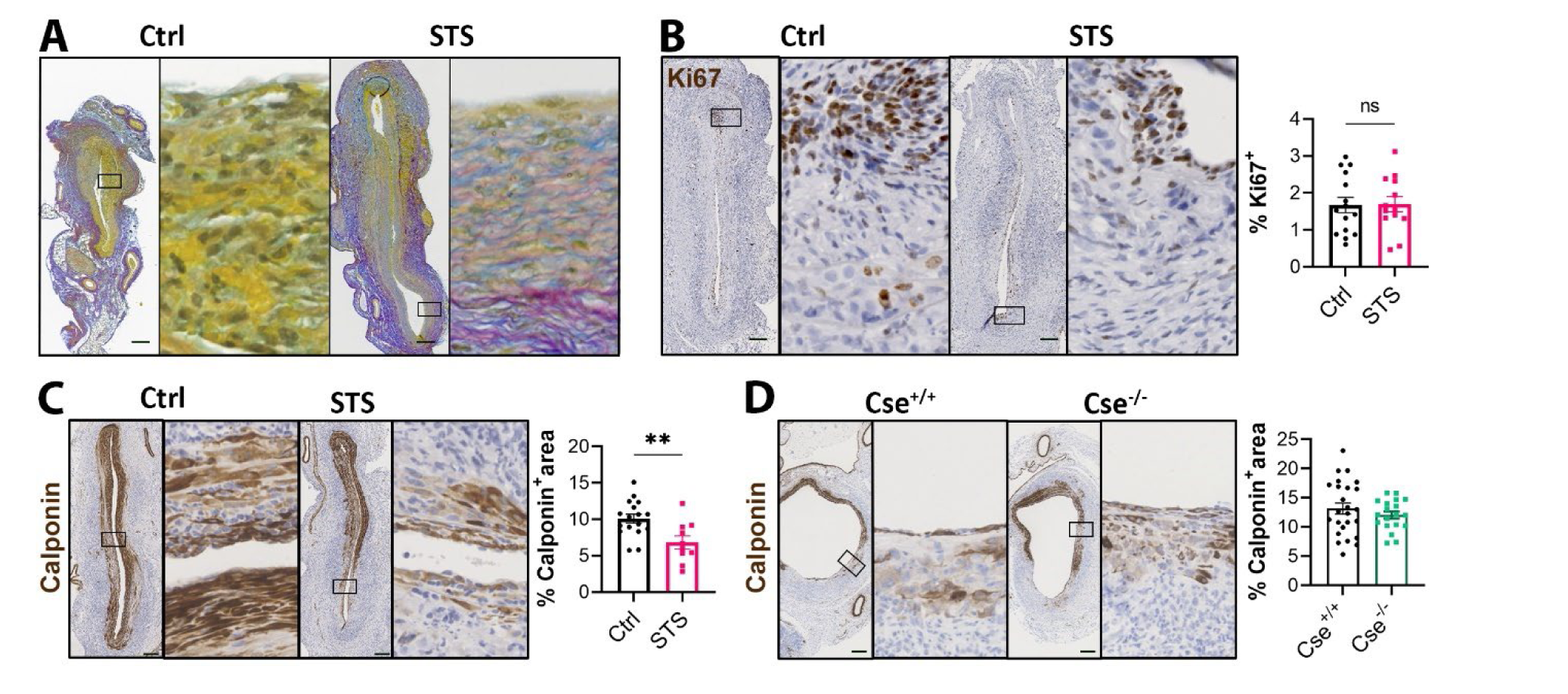
STS reduces VSMC coverage in AAA. Representative Herovici staining (A), and Ki67 (B) and Calponin (C) immunostaining in AAA section from WT mice treated or not (Ctrl) with STS. D) Representative Calponin immunostaining in sub-renal AAA in Cse^+/+^ and Cse^-/-^ mice. Data are mean±SEM of 10 to 18 animals per group. Scale bar 100 μm. Right insets are 5-fold magnification of main image. **p<0.01 as determined by bilateral unpaired t-test.

**Figure 6.**
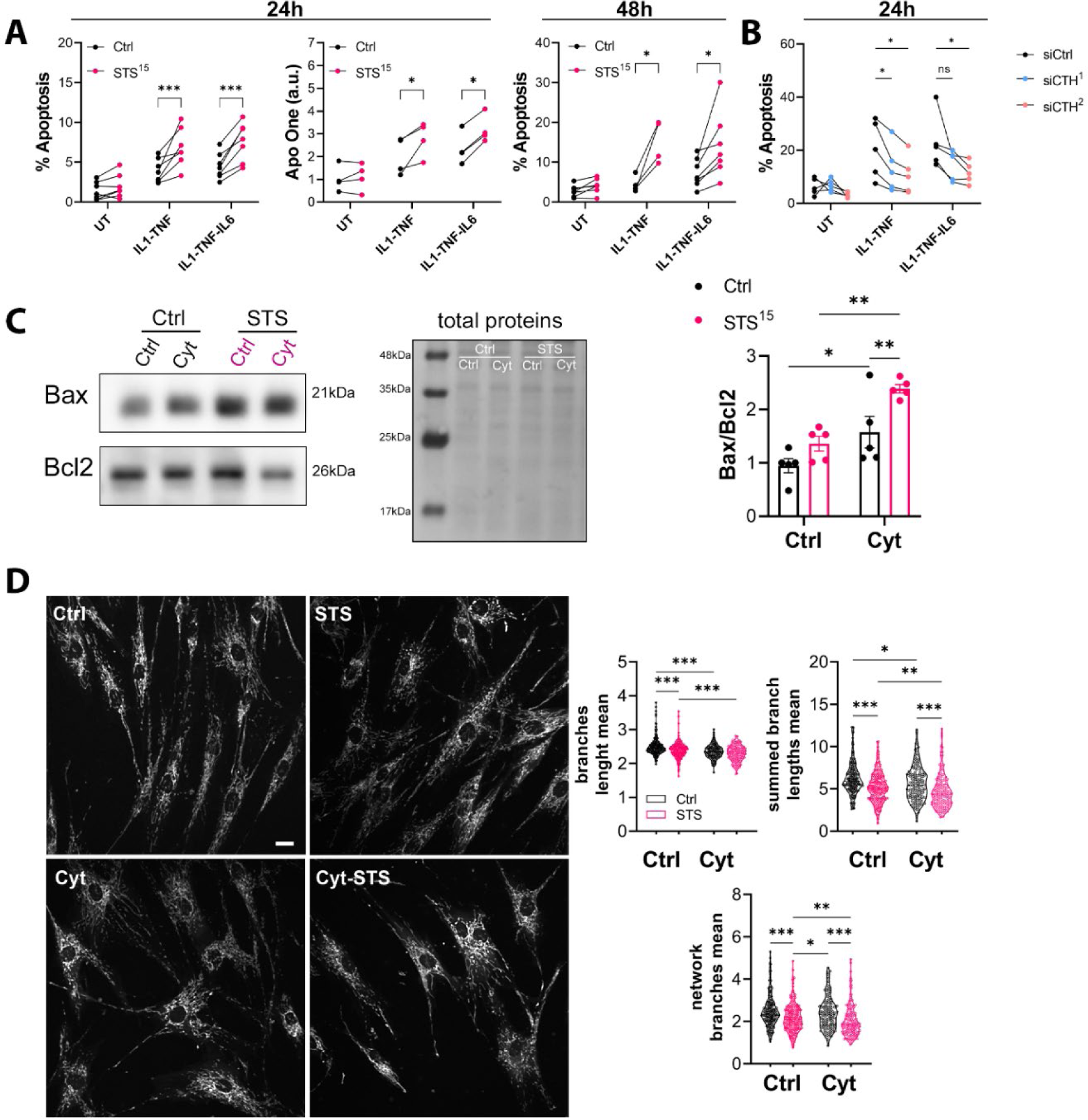
STS increases cytokine-induced mitochondrial dysfunction and VSMC apoptosis. A) Apoptosis or cleaved caspase 3 activity (Apo One) in VSMC treated or not (UT) for 24 h or 48h with cytokines or 15mM STS, as indicated. B) Apoptosis in VSMC knocked down for CTH using 2 distinct siRNAs (siCTH ^1^ and ^2^), and treated or not with cytokines as indicated. C) Representative western blot and quantitative assessment of Bax over Bcl2 protein levels in VSMCs treated or not with a mix of cytokines IL-1β+TNFα+IL-6 (Cyt) and/or 15mM STS for 24h. A-C) Data are mean±SEM of 4 to 6 independent experiments. *p<0.05, **p<0.01, ***p<0.001 as determined by Mixed-effects model (REML) with Šídák’s multiple comparisons tests. D) Representative images and quantitative assessment of live Mitotracker staining in VSMC treated with IL-1β+TNFα+IL-6 (Cyt) and/or 15mM STS. Data are mean±SEM of 5 independent experiments. *p<0.05, **p<0.01, ***p<0.001 as determined by Kruskal-Wallis tests corrected by Dunn’s.

## Discussion

Given the anti-inflammatory and antioxidant properties of H_2_S^7, 13^, we hypothesized that STS, a clinically relevant source of H_2_S^14, 15^, would protect against aneurysm growth in a mouse model of elastase-induced AAA. Surprisingly, STS promoted AAA growth and rupture, whereas Cse^-/-^ mice were protected. AAA’s primary pathogenic features are i) infiltration of innate and adaptive immune cells in the aorta wall; ii) proteolysis of the extracellular matrix (ECM), and iii) loss of VSMC^5, 37, 38, 40^. Here, we show that STS inhibits the infiltration of innate and adaptive immune cells, but facilitates VSMC apoptosis and proteolysis of the ECM, overall leading to increased AAA growth and rupture.

Inflammation is a key hallmark of AAA pathology^5, 40^, involving the infiltration of immune cells like neutrophils, macrophages, and CD86^+^ antigen-presenting cells (APC) involved in regulating immune responses via promotion of T cell activation and cytokine production. As expected for a H_2_S donor, STS reduces immune cell infiltration, including macrophages, CD86^+^ APC and lymphocytes. Conversely, loss of CSE increased infiltration of macrophages, CD86^+^ APC and lymphocytes. Neutrophil recruitment was not impacted by STS or Cse, but 14 days post-op might be too late to accurately measure neutrophil in that model^19^. This data suggest that CSE/H_2_S impact the innate immune response, which extend to reduced adaptive immune response as well. Despite this anti-inflammatory effect, H_2_S has a negative impact on AAA growth is our model. This can be explained in many ways.

First, the anti-inflammatory effect of STS could be detrimental. Indeed, all immune cell populations have been described in AAA, with different types promoting or limiting AAA growth^5, 40^. Here, STS had a general anti-inflammatory effect, reducing the infiltration of all macrophages, including CD206^+^ M2 macrophages, which have been shown to promote vascular repair ^40, 41^. Other immune cells with beneficial effects such as CD4^+^ regulatory T cells might be similarly impacted, leading to AAA growth. Our findings contradict studies that demonstrate H_2_S can enhance M2 macrophage polarization^42^, but they do support other research that show exogenous H_2_S (NaHS) suppresses both M1 and M2 macrophage invasion^43, 44^. Further studies are required to decipher the exact effect of H2S on sub-populations of immunes cells in the context of AAA.

Second, STS may facilitate ECM degradation. The pathogenesis of AAA is characterized by a breakdown of elastic and collagen fibers due to increased proteolytic activity, mainly by MMP-1, −2, and −9^37, 38^. According to our proteomics data, STS lowers the protein levels of elastin and a few collagen proteins. In our model, STS/CSE also increases MMP9, which may accelerate ECM degradation. In this elastase-induced AAA model, H_2_S likely enhances ECM proteolysis. Although STS has a minor effect on ECM proteins, this could contribute to AAA expansion, especially since Elastin appears to be among the primary targets of STS.

Third, STS inhibits VSMC proliferation. The VSMC maintain and renew the ECM and ensure the structural integrity of the aortic wall. VSMC dedifferentiation in synthetic cells secreting a large quantity of matrix remodeling proteins contributes to the progression of AAA^37, 38^. Phenotypic modulation takes place early in aneurysm development in both human aneurysm samples and mouse models, and results in VSMC clonal expansion to compensate loss of structural integrity {Clement, 2019 #1552}. Despite pre-clinical studies using various mouse models showing that anti-inflammatory medications prevent AAA formation and/or dissection, anti-inflammatory medications in humans promote AAA growth. This adverse outcome is most likely caused by the cytostatic effect of immunosuppressive medications on VSMC^45, 46^, showing that VSMC expansion can hinder AAA growth. These synthetic VSMC lack the expression of specific VSMC markers such as Calponin^37, 38^. In our experiment, STS reduced Calponin^+^ cell coverage but does not appear to affect the phenotypic of VSMC, implying that the reduced staining result of a reduced number of VSMC. Previously, we and others demonstrated that CSE and other H_2_S donors, including STS, suppress VSMC proliferation^14, 23, 47–49^. According to the Ki67 staining, STS in this case did not inhibit cell proliferation in the aortic wall. However, the Ki67 staining did not make it possible to differentiate between the proliferation of VSMC and other cells, especially immune cells. In addition, STS may impact VSMC proliferation at an earlier time point in the model of elastase-induced AAA. We believe that the effect of STS on VSMC proliferation plays a significant role in AAA growth by preventing VSMC expansion to stabilize the weakened aortic wall in the early phase of AAA formation post-surgery^41^. Further studies are required to test this hypothesis.

Last, STS increases cytokine-induced loss of VSMC. VSMC apoptosis is a hallmark aortic aneurysms progression and rupture^50^. Our analysis of cleaved caspase 3 in the aortic wall did not allow us to evaluate apoptosis *in vivo*. This is probably due to the rapid phagocytosis of dying cells by macrophages *in vivo*. *In vitro* findings revealed that STS alone is not cytotoxic, as previously demonstrated^14^, but facilitates cytokine-induced VSMC apoptosis. This is not surprising, as elevated levels of exogenous H_2_S may cause cell cycle arrest and apoptotic cell death^35, 50–52^. Similarly to our findings, H_2_S has been previously showed to promote ROS-induced mitochondrial apoptosis via the Bax/Bcl2^35, 53^. STS-induced cell cycle arrest probably tips the balance between pro and antiapoptotic signals toward apoptosis even though STS may induce cytoprotective mechanism. Indeed, we observed that STS promotes mitochondrial biogenesis, which is associated with improved function, stress resistance and cell survival^35, 54, 55^. STS stimulates the expression of the master regulator of mitochondrial biogenesis PGC-1α and downstream transcription factor NRF1. This is most likely owing to a minor inhibitory effect on mitochondrial respiration and ATP production^14^. This, in turn, increases AMPK activity, Sirtuins, and increased PCG-1α expression, leading to increased mitochondrial components as observed in our proteomic analysis. Supporting this hypothesis, glycolysis and fatty acid oxidation were also up in our proteome analysis, probably because of reduced oxidative phosphorylation. This was observed in native aorta, and it is unlikely that STS stimulates mitochondrial biogenesis in the pro-inflammatory environment of the AAA.

Overall, we propose that STS has both beneficial and deleterious effects in the context of AAA. Thus, STS has anti-inflammatory properties and promotes mitochondrial biogenesis. However, STS also reduces VSMC proliferation, facilitates ECM degradation and VSMC apoptosis in a pro-inflammatory environment, leading to AAA growth.

### Limitations

The main limitation of our study is the dose of STS (4g/L; 0.5g/Kg/day) used in our experiments. The dosage of STS used in this study is comparable to previous experimental studies using oral administration at 0.5 to 2 g/Kg/day^58–61^, and we recently showed that this dose of oral STS is not toxic in mice^14^. However, we also observed in another study that 2g/L of STS was more potent in stimulating revascularization than 4g/L, suggesting a very narrow therapeutic range for STS^15^. It is well known that H_2_S may exhibit cytotoxic effects^62^ and our study indicates that STS may facilitate cytokine-induced apoptosis and AAA growth. Lower dose of STS should be tested as it may alleviate the negative effect of STS on VSMC. However, it might also lessen the anti-inflammatory effect of STS. Of note, our findings also document that CSE, hence endogenous H_2_S production, is also deleterious in the model of elastase-induced AAA. Therefore, the negative impact of STS on AAA growth is not due solely to the dose of STS used in our study. Cse^-/-^ mice might develop smaller AAA due to reduced MMP9 and matrix remodelling. Although we did not observe an increased VSMC coverage in Cse^-/-^ mice, loss of Cse might also facilitate positive wall remodelling as CSE inhibits VSMC proliferation and migration^14, 23, 47–49^.

Our results are in contradiction with several studies reporting beneficial effects of Cse and H_2_S donors against the formation of aortic aneurysm and dissection^9, 11, 12^. Of note, these previous studies all employed various models of angiotensin II-induced aortic dissection. In these models, vasoreactivity plays a major role to counterbalance angiotensin II-induced vasoconstriction and hypertension. H_2_S is a commonly known vasodilator^63^, and H_2_S donors, including STS, have been reported to lower blood pressure in various models of hypertension^64, 65^, including angiotensin II-induced hypertension^58, 66^. The vasoactive property of Cse/H_2_S likely provided additional protection against aortic dissection is these models. Thus, Zhu et al. demonstrated that Cse^-/-^ mice are more sensitive to angiotensin II-induced aortic elastolysis and medial degeneration and are rescued by NaHS treatment^9^. It should be noted that these Cse^-/-^ mice, developed by Prof. Rui Wang, are hypertensive^67^, and that the NaHS treatment normalizes blood pressure in this study, event when treated with angiotensin II^9^. Our Cse^-/-^ mice, developed in collaboration with late Prof. James R. Mitchell, are normotensive^14^, like the Cse^-/-^ mice developed by Prof Isao Ishii^68^. That said, in line with previous studies, we also observed increased elastin breaks in Cse^-/-^ mice, leading to aortic dissection although our model of periadventitial elastase is not reported to induce aortic dissection^20^. This aortic wall dissection did not lead to aortic rupture, but may indicate increased aortic stiffness related to impaired vasoreactivity. Conversely, the vasorelaxant property of STS may facilitate AAA growth in the elastase-induced AAA model.

In recent years, the angiotensin II model has been used extensively for aneurysm research. However, the major mechanism of Angiotensin II-induced AAA in ApoE^-/-^ mice is subsequent to aortic dissection, which is distinct from human AAA. Combination of BAPN with Ang II increased the incidence of aneurysm in WT mice. However, dissections and ruptures in those models all occur within the first week during the acute phase of aneurysm induction. Aortic aneurysms in the Angiotensin II model also develop in the suprarenal aortic segment, whereas 70% of human aneurysm occur in the infrarenal section. The elastase-induced AAA model is regarded as the best model for human AAA disease^16, 17^. A recent single-cell RNA analysis comparing various aneurysm models revealed that the elastase model shows the closest signature to human AAA^69^. Therefore, our data provided much needed perspective into the impact of CSE and H_2_S on the aortic wall in normotensive conditions. Given the inherent bias of working with a vasodilating agent, we believe models of Angiotensin II-induced aortic dissection should be avoided. Further research employing different AAA models, such as the CaCl_2_ model^16, 17^, might be beneficial in better understanding the role of H_2_S in AAA formation.

## Conclusion

In summary, in our experimental conditions, STS, a clinically authorized substance, decreases inflammation but has a detrimental impact on vascular remodelling and AAA formation. The significance of the adverse impact of STS on the advancement of AAA is of importance considering the increasing utilization of STS in clinical settings for a variety of indications. STS is already used for the treatment of acute calciphylaxis, a rare vascular complication of patients with end-stage renal disease^70, 71^. STS is also tested in a several clinical trials for the treatment of ectopic calcification (NCT03639779; NCT04251832; NCT02538939), or to reduce myocardial infarct size in ST-segment elevation myocardial infarction (STEMI) patients with percutaneous coronary intervention (NCT02899364). Our finding calls for randomized controlled trials testing long-term administration of STS to further explore the safety and effects of STS administration on the vascular wall.

Our findings revealed that H_2_S effectively attenuates inflammation but does not impede the development of AAA. This study provides evidence of H_2_S inefficacy in mitigating AAA growth and identifies a deleterious influence of H_2_S on VSMC in this context, highlighting the complex role of H_2_S in AAA progression.

## Author Contributions

Conceptualization, F.A. and S.D.; methodology, F.A., S.D, D.M., M.L. and S.U.; validation, F.A., S.D.; formal analysis, D.M., C.B., F.A., S.U. and M.L; investigation, D.M., C.B., F.A., S.U. and M.L.; writing—original draft preparation, C.B. and F.A.; writing—review and editing, F.A., C.B., M.L. and S.D.; visualization, C.B. and F.A.; supervision, F.A. and S.D.; project administration, F.A. and S.D.; funding acquisition, F.A. and S.D. All authors have read and agreed to the published version of the manuscript.

## Data Availability Statement

The data presented in this study are available on request from the corresponding author.

## Acknowledgments

We thank the Mouse Pathology Facility of the Faculty of Biology and Medicine, University of Lausanne, Lausanne, Switzerland for their services in histology. We thank the Cellular Imaging Facility of the Faculty of Biology and Medicine, University of Lausanne, Lausanne, Switzerland for their services in microscopy. We thank the Protein Analysis Facility of the Faculty of Biology and Medicine, University of Lausanne, Lausanne, Switzerland for the Mass spectrometry-based proteomics work.

