## supplementary data and figures for "The hydrogen sulfide donor sodium thiosulfate limits inflammation but aggravate smooth muscle cells apoptosis and aneurysm progression in a mouse model of abdominal aortic aneurysm"

**Running title:** sodium thiosulfate promotes AAA growth.

**Category**: original article

**Keywords**: abdominal aortic aneurysm; AAA; hydrogen sulfide; $H_2S$; thiosulfate; vascular smooth muscle cells; elastase.

***The following are supplementary data related to this study***

**Supplementary Table S1: Antibodies**

| Target antigen | Vendor | Catalog # | Working concentration |
|---|---|---|---|
| **Caspase 3** | Cell signaling | #14220T | 1/300 |
| **SMA** | Cell signaling | #19245S | 1/500 |
| **MPO** | Abcam | ab208670 | 1/2000 |
| **CD86** | Cell signaling | #195895 | 1/1000 |
| **MMP9** | Abcam | ab228402 | 1/500 |
| **CD3** | Cell signaling | #989415 | 1/100 |
| **CD8** | Cell signaling | #999405 | 1/100 |
| **CD206** | Abcam | ab64693 | 1/20 000 |
| **Calponin** | Abcam | ab46794 | 1/100 (IHC-P)<br>1/5000 (WB) |
| **Ki67** | Abcam | ab16667 | 1/200 |
| **F4/80** | Invitrogen | MF48000 | 1/50 |
| **Bax** | Santa Cruz | Sc- 526 | 1/200 |
| **Bcl2** | Santa Cruz | Sc- 492 | 1/200 |
| **SM22α** | Abcam | ab14106 | 1/20 000 |
| **α-SMA** | Cell signaling | #19245 | 1/1000 |
| **Oxphox cocktail** | Abcam | ab110413 | 1/1000 |
| **Cgl** | Proteintech | 12217-1-AP | 1/1000 |

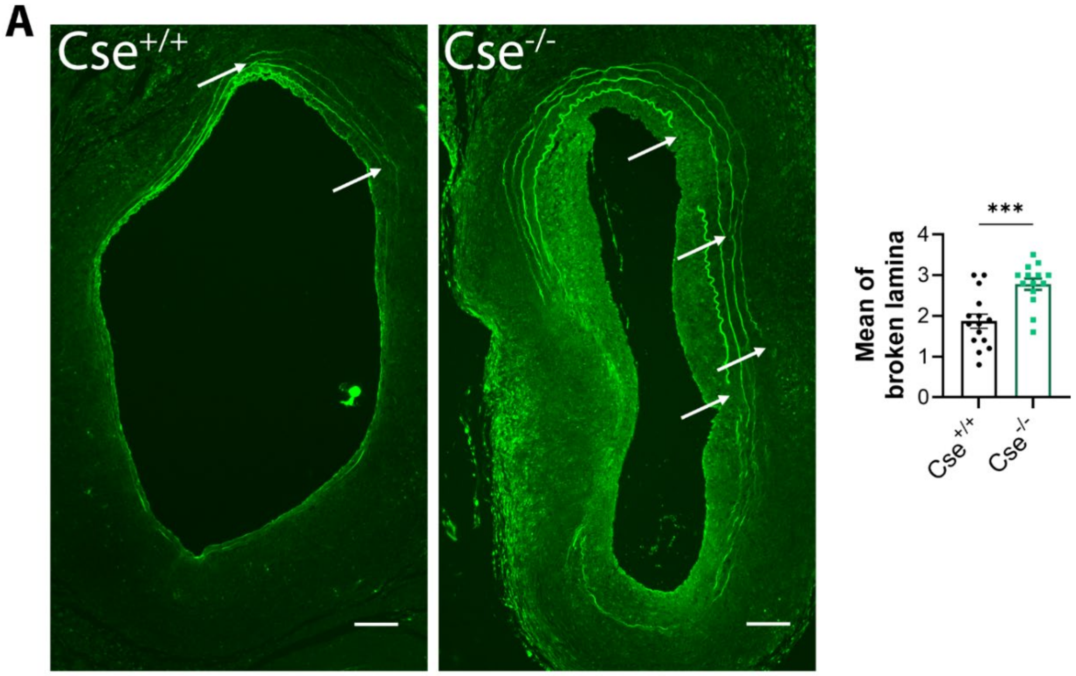


Figure S1. Cse$^{-/-}$ display increased incidence of elastin breaks.

Representative autofluorescence AAA of Cse$^{+/+}$ and Cse$^{-/-}$ mice. White arrows show broken lamina. Data are mean±SEM of 15 to 18 animals per group. ***p<0.001 as determined by bilateral unpaired t-test. Scale bar 100 µm.

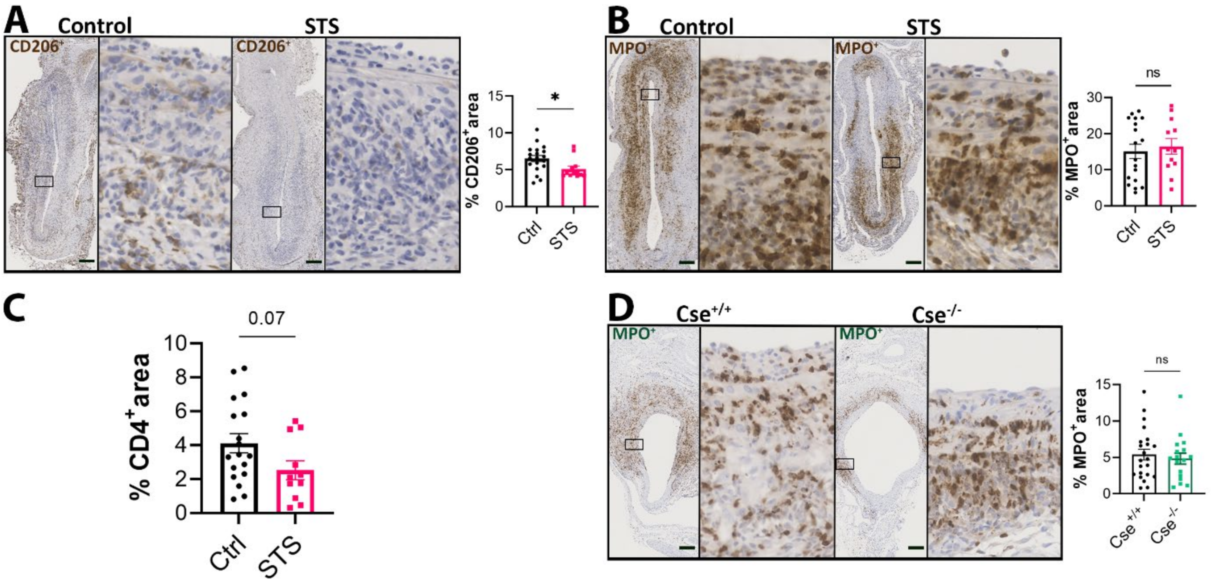


Figure S2. STS decreases inflammation in the aortic wall

Representative CD206 (A), MPO (B) immunostaining in sub-renal mouse aorta in WT mice with topical elastase application, treated or not (control) with 4g/L STS (STS). Quantification of CD4+ cells in AAA (C). Representative MPO (D) immunostaining in sub-renal mouse aorta in Cse+/+ and Cse-/- with topical elastase application. Boxes next to AAA are 5-fold magnifications of AAA. Data are mean±SEM of 15 to 18 animals per group. Scale bar 100 µm. *p<0.05 as determined by bilateral unpaired t-test.

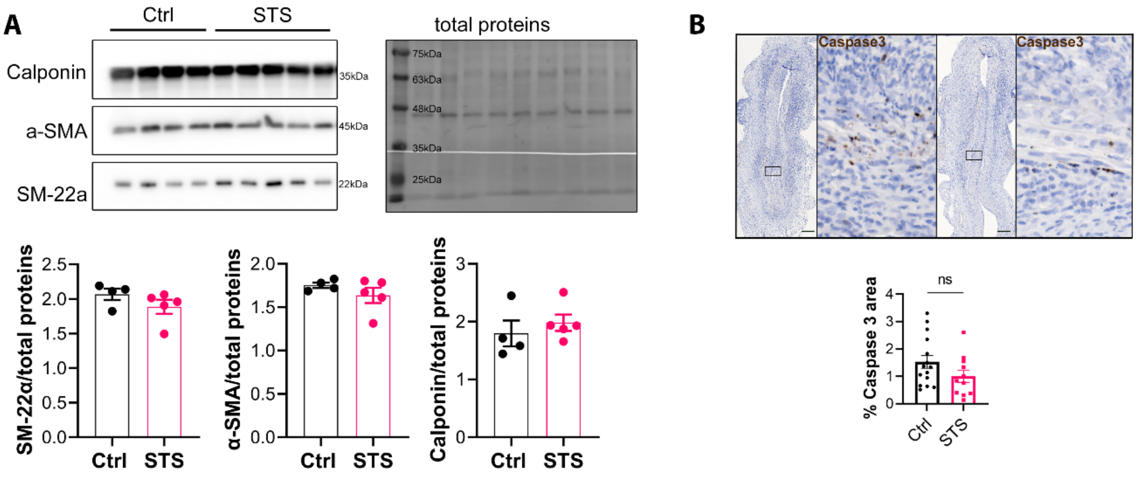


Figure S3. STS did not impact VSMC phenotype

A) Western blot analysis in native aorta from mice treated or not (Ctrl) for 1 week with STS. Data are mean±SEM of 4 mice per group. No significant difference as determined by bilateral unpaired t-test. B) Representative Caspase 3 immunostaining in sub-renal mouse AAA in WT mice with topical elastase application treated with STS (STS) or not (Ctrl). Right insets are 5-fold magnifications of left images. Scale bar 100 µm. Data are mean±SEM of 15 to 18 animals per group. No significant difference as determined by bilateral unpaired t-test.

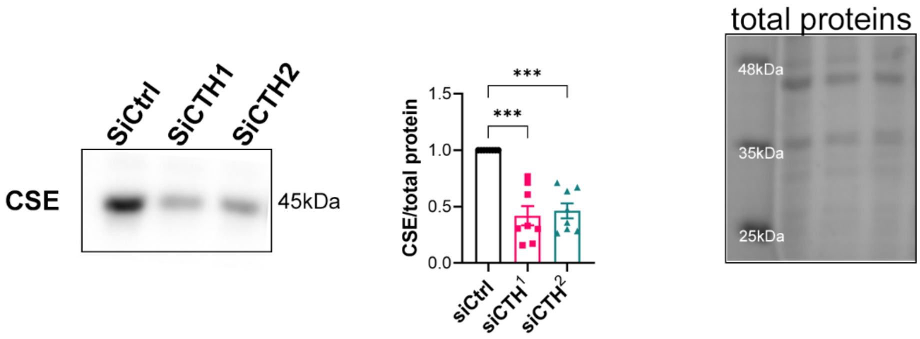


Figure S4. siRNA-mediated CTH knock-down in VSMC

Western blot analysis of CSE protein in VSMC transfected with a control siRNA (siCtrl) or two distinct siRNA targeting CTH (siCTH [1] and [2]). ***p<0.001 as determined by repeated measures One-way ANOVA followed by Dunnett's multiple comparisons tests.
